# *In-silico* identification of potential antigenic proteins in *Bartonella bacilliformis* for the serological diagnosis of Carrions’ disease

**DOI:** 10.1101/2022.06.06.494961

**Authors:** Victor Jimenez-Vasquez, Karen Calvay-Sanchez, Yanina Zarate-Sulca, Joaquim Ruiz, Giovanna Mendoza-Mujica

## Abstract

The current methods for inferring antigenic, pathogenic or virulence factors in bacteria include *in-silico* or computational tools. *Bartonella bacilliformis*, the causal agent of Carriońs disease (CD) is a Gram-negative microorganism transmitted by the sand fly *Lutzomya verrucarum* mainly in Peruvian Inter-Andean valleys. A better understanding of the pathogenicity of *B. bacilliformis*, the implementation of serological diagnostic methods and the development of candidate vaccines for the control of CD could be facilitated by the identification of outer membrane exposed proteins such as outer-membrane beta-barrels (OMBB) and outer-membrane lipoproteins (OMLPP). In this study we present the *in-silico* identification of OMBBs and OMLPPs in *B. bacilliformis*. The present analysis identified 32 OMBBs and 9 OMLPPs, of which 15 and 4 are reported for the first time as potential antigenic, pathogenic and virulence-related proteins, respectively. Future implications of this study are discussed and compared with proteins validated by experimental assays.

## Introduction

Carrion’s disease (CD) is an endemic illness of Andean countries such as Peru and Ecuador, with an additional historical presence in Colombia [1, 2]. In Peru, CD represents one of the main challenges for public health due to poverty and poor sanitation in endemic localities, affecting children with chronic malnutrition. *Bartonella bacilliformis*, the etiological agent of CD, is a Gram-negative pleomorphic microorganism transmitted by sand flies of the *Lutzomyia* genus, especially *L. verrucarum* [2]. Climate change and variability in inter-Andean valley rainfall associated with the El Niño-Southern oscillation (ENSO) favors the reproduction of L. verrucarum and the consequent emergence of local CD outbreaks with a non-negligible number of cases [3]. CD clinical manifestations are diverse as the microorganism parasites human erythrocytes generating an “acute phase, called Oroya fever, characterized by both anemia and febrile illness [2]. Nevertheless, the nature of the initial symptoms may be confused with that of other infectious diseases, such as malaria, dengue or others. This is of special relevance because the absence or delay of adequate treatment may result in fatal outcomes [2]. In this sense, mortality rates among untreated or inadequately treated patients of up to 88% have been described [4]. Of note, in Peru, the overall Oroya fever lethality ranges between 0.5 and 3%, with about 10% of severe cases attending reference hospitals having a fatal outcome [2].

Eventually, after a period of several weeks to months following the acute phase, a non-life threating eruptive or “verrucose phase” distinguished by notable skin eruptions develops and may be presented in the absence of a previous acute infection [1, 2].

In addition to the clinical manifestations described above, patients with asymptomatic bacteremia have been identified and are considered as potential reservoirs of *Bartonella bacilliformis* and a potential source of infection for susceptible persons [3].

While bacteriological cultures are accurate, they are time-consuming given the slow growth rate of *B. bacilliformis* (about 4-15 days), and therefore have limited usefulness for diagnostic purposes. On the other hand, blood smear detection is fast, accurate and low cost, and is thereby largely used for diagnostic purposes, especially in rural areas. Nevertheless, it has a low sensitivity, generating false negatives, limiting the confirmation of any diagnosis [5, 6]. Molecular methods based on the detection of specific gene regions such as gltA, ribC and ialB have been proposed, being either limited to gene amplification or combined with a subsequent process, such as enzymatic digestion [7]. Notwithstanding, in addition to the difficulties in implementing these techniques in most rural areas affected, some of these genes are not specific for *Bartonella bacilliformis*, causing cross reactivity with other pathogens [7]. It should be taken into account that the current serological techniques available are in-house with no commercial development, having a limited specificity, and potential cross reactivity with other *Bartonella*ceae or other microorganisms cannot be ruled out, similar to what has been reported for different immunological approaches for the detection of *Bartonella* quintana and *Bartonella* henselae infections [8, 9]. Regarding rapid diagnostic tools that can be used in endemic areas, different antigenic candidates have been proposed but none has been introduced in clinical practice [10, 11].

Data about immune response to *B. bacilliformis* is limited, with antibody immunity build-up for the development of partial immunity being described considering immunoglobulin M (IgM) as a biomarker of the acute phase and immunoglobulin G (IgG) as a marker of previous exposure [10, 12].

The finding of specific outer membrane proteins (OMPs) of *B. bacilliformis* could establish the basis for the future implementation of an accurate and sensitive serological assay for the diagnosis of CD as OMPs activate immune response and participate in virulence mediating pathogen-host interactions [13]. In Gram-negative bacteria, the tertiary structure of OMPs includes beta-barrel structure composed of a variable number of beta-strands. Another type of exposed OMPs known as outer membrane lipoproteins are also involved [14, 15]. In addition, both outer-membrane beta-barrels (OMBB) and outer-membrane lipoproteins (OMLP) could be good candidates for the development of vaccines, immunological target tests and could promote better understanding of the pathogenicity of *B. bacilliformis*, facilitating the identification of therapeutic molecules for *in-silico* and experimental assays [16].

Identification of the antigenic-determinant is crucial for designing immune treatment such as a vaccine against infectious diseases, or to avoid cross-reactivity of antibodies used in diagnostic methods [17]. Immunoinformatics addresses the molecular interactions of potential binding sites by computational methods [18] and is the best alternative to recognize epitopes.

Immunoinformatics is inexpensive, accurate and not time-consuming, allowing the design and synthesis of a molecule to replace the antigen [17].

The present study aimed to identify exclusive potential antigenic candidates of *B. bacilliformis* represented by OMBB and OMLP.

## Materials and Methods

### Data preparation

Potential antigenic proteins of *B. bacilliformis* based on the complete genome of the strain KC584 (GenBank code CP045671.1) were identified. Functional genes (CDSs) were downloaded in fasta format and all headers were edited in order to retain the name, orientation and position in the genome. Nucleotide sequences were translated with VirtualRibsome 2.0 [19] (https://services.healthtech.dtu.dk/service.php?VirtualRibosome-2.0) and peptide sequences were downloaded in fasta format.

### Identification of lipoproteins, beta-barrel and experimental proteins

Outer membrane lipoproteins, outer membrane beta-barrel proteins and experimental proteins were identified with antigenic re-activity by using different approaches. Thus, signal peptides were detected with PrediSi [20] (http://www.predisi.de/), SignalP 5.0 [21] (http://www.cbs.dtu.dk/services/SignalP/) and Phobius [22, 23] (https://phobius.sbc.su.se/) predictors. Lipoproteins were identified with SignalP-5.0, LipoP 1.0 [24] (http://www.cbs.dtu.dk/services/LipoP/), LipoCBU [25] (http://services.cbu.uib.no/tools/lipo), SpLip [26] (http://www.iq.usp.br/setubal/tools.html) and PredLipo [27] (http://bioinformatics.biol.uoa.gr/PRED-LIPO/input.jsp) predictors. Beta-barrel proteins were identified with mcmbb [28] (http://athina.biol.uoa.gr/bioinformatics/mcmbb/), BOMP [29] (http://services.cbu.uib.no/tools/bomp), OMPdb [30] (www.ompdb.org), PredTMBB2 [31] (http://www.compgen.org/tools/PRED-TMBB2) predictors. Subcellular localizations were inferred with Cello [32] (http://cello.life.nctu.edu.tw/cello2go/), LocTree3 [33] (https://rostlab.org/services/loctree3/), Psort-B 3.0 [34] (https://www.psort.org/psortb/), Gneg.mPloc [35] (http://www.csbio.sjtu.edu.cn/bioinf/Gneg-multi/). Furthermore, 9 proteins with antigenic re-activity based on scientific studies were found in the PubMed repository.

Lipoproteins with the following three characteristics were selected: 1) containing signal-peptide potentially cleaved by signal peptidase II (SPaseII) in 3 out of the 5 tools, 2) located on the outer membrane in 3 out of 4 tools, and 3) recognized as lipoproteins by 3 out of the 5 tools. Beta-barrel proteins with the following two characteristics were selected: 1) located in the outer membrane in 3 out of 4 tools and 2) recognized as beta-barrel by 3 out of 4 predictive tools.

### Comparison of selected proteins

In order to retain only *B. bacilliformis*-exclusive-proteins, parallel comparisons of selected proteins were conducted with proteins belonging to *Mus musculus* and *Homo sapiens* with a Bit-score < 100, % identity < 35, % coverage < 35; febrile illness-associated bacterial species belonging to genera *Anaplasma*, *Brucella*, *Ehrlichia*, *Leptospira*, *NeoRickettsia*, *Orientia* and *Rickettsia* with a Bit-score < 100, 43% < identity < 95% and 43% < coverage < 95%, and *Bartonella* genus with a Bit-score < 100, 43% < identity < 95%, 43% < coverage < 95 %. To perform this step, we used the BLAST+ tool. Thus, the selected *B. bacilliformis* proteins were compared with proteomes derived from genomes with a minimal coverage depth of 100x. The workflow is shown in Figure 1.

**Figure 1.**
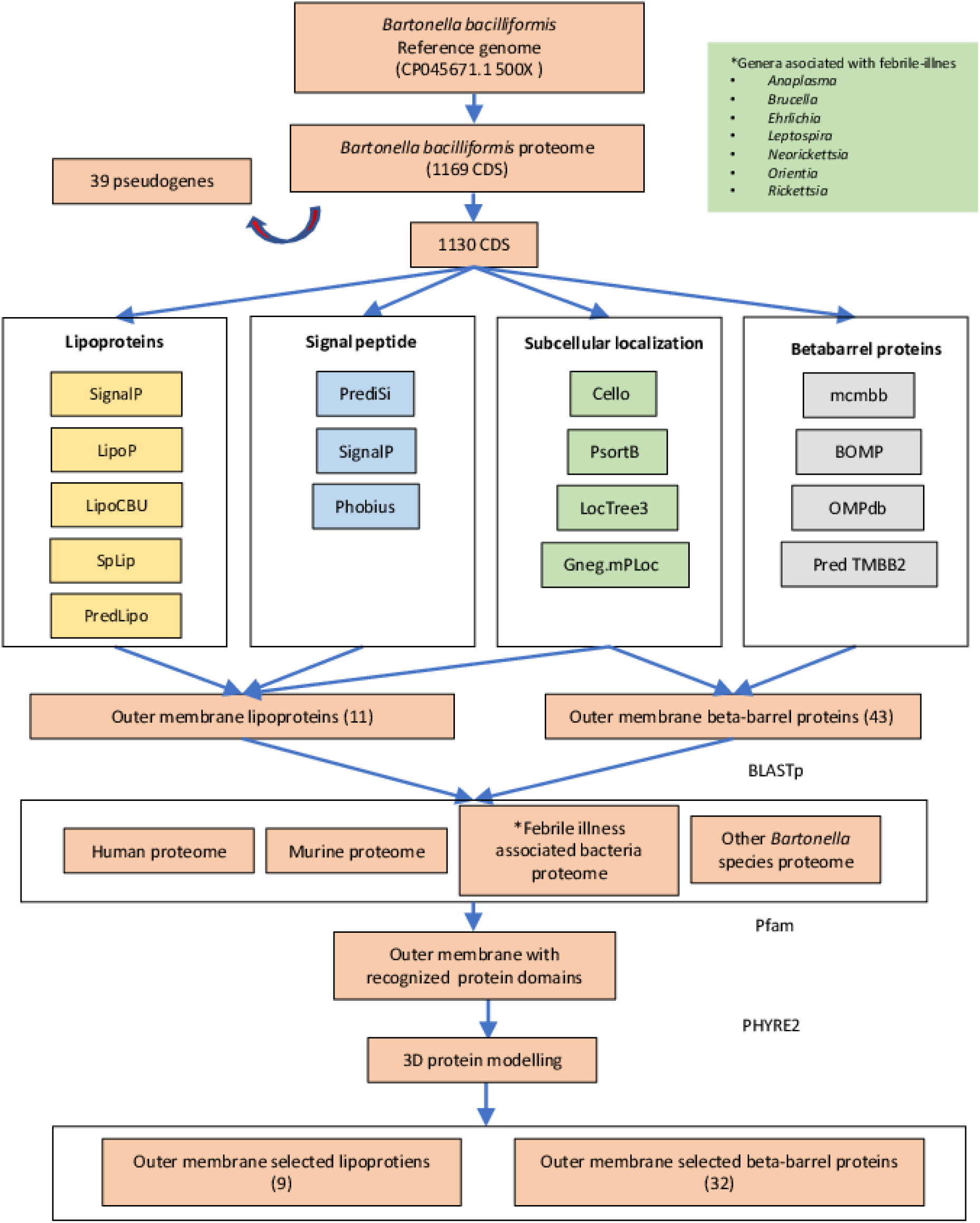
Workflow.

### 3D-mapping

A series of 3D structures were obtained with the Phyre2 online program [36] (http://www.sbg.bio.ic.ac.uk/phyre2/). We identified protein domains with the PFam program and checked models in Chimera.

## Results

The present study aimed at finding potential antigenic OMPs and the identification of potential antigenic domains in proteins with known immunological reactivity in *B. bacilliformis*. By contrasting subcellular localizations, the presence of signal peptides, lipoprotein and beta-barrel protein predictors followed by proteome-proteome comparisons between human, murine, febrile-associated and other *Bartonella* species, 32 OMBB proteins and 11 OMLPs were identified, which are shown in Tables 1 and 2. All 9 experimental proteins with immunological reactivity followed the same procedure (Table 3). For the present descriptions, selected proteins were listed in terms of accession numbers followed by their order in the annotated genome (GenBank accession: NZ_CP045671.1).

**Table 1.**
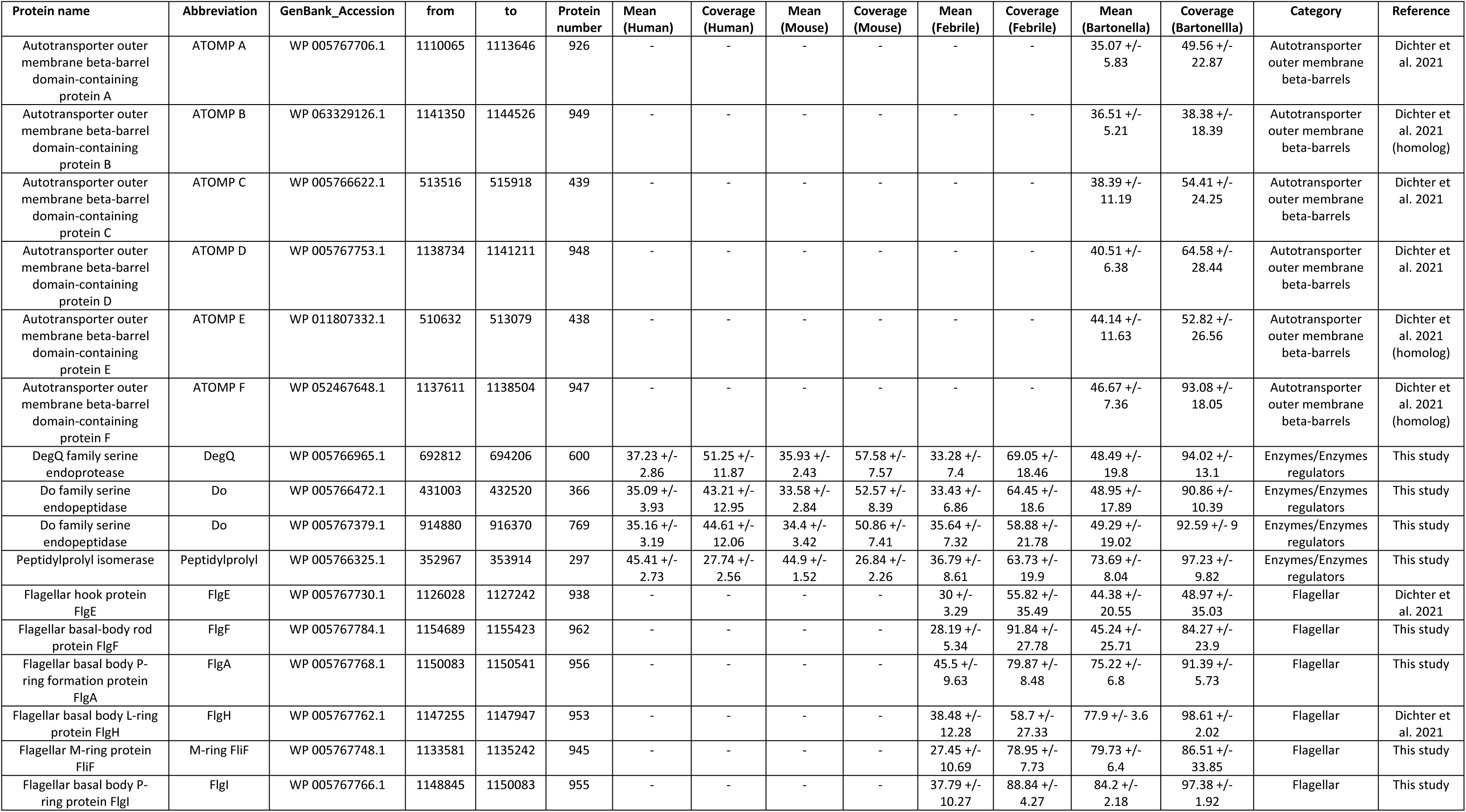
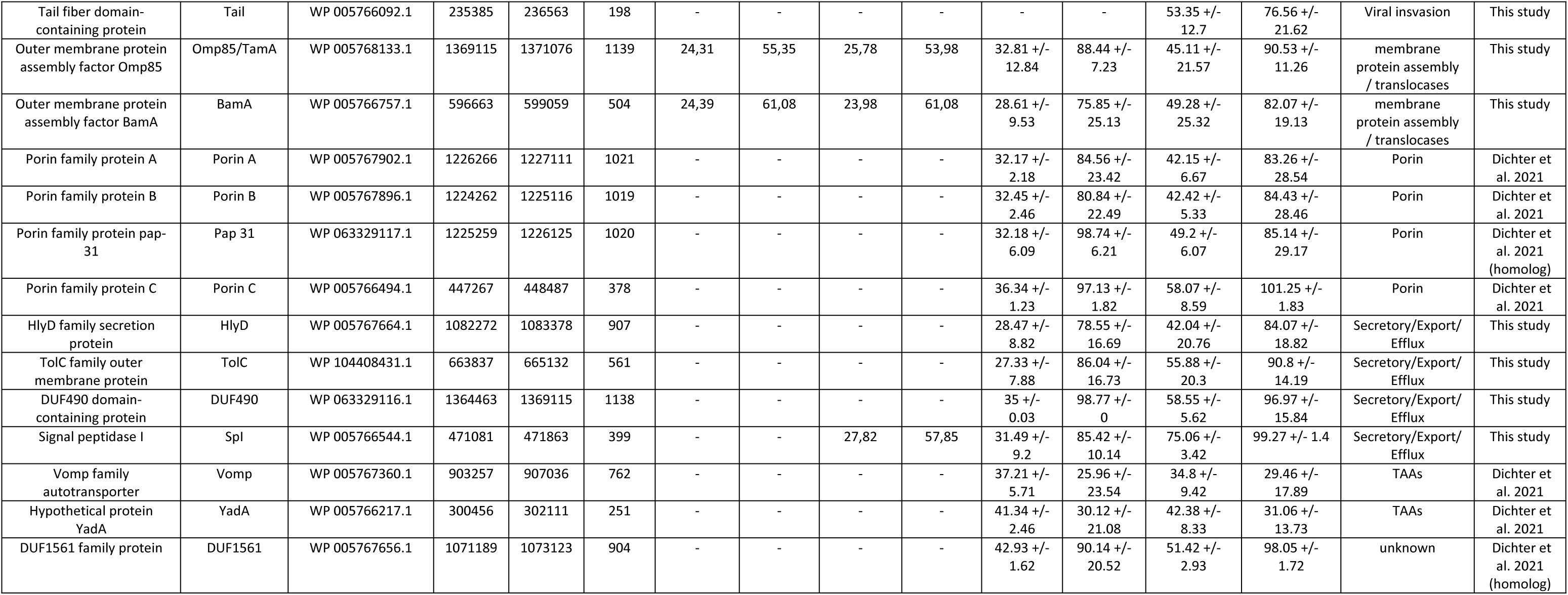
Outer membrane beta-barrel proteins identified in this study.

**Table 2.**
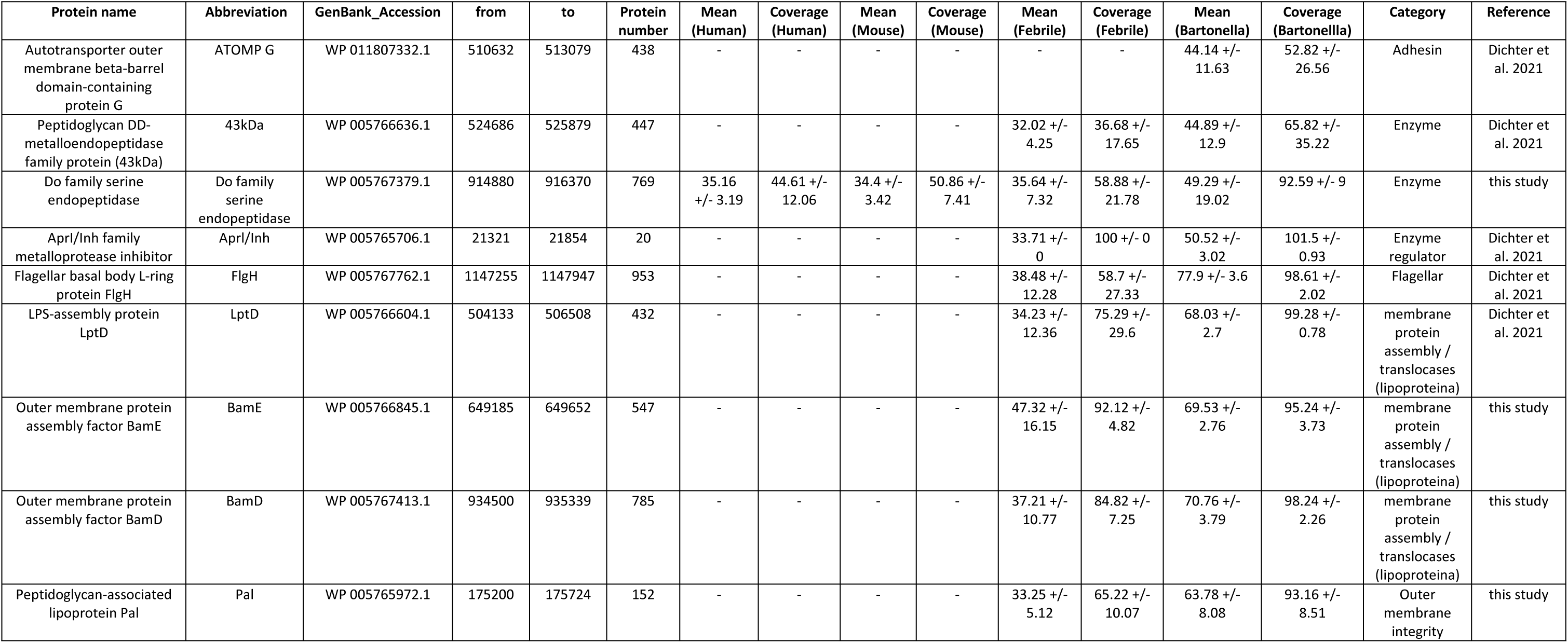
Outer membrane lipoproteins identified in this study.

**Table 3.**
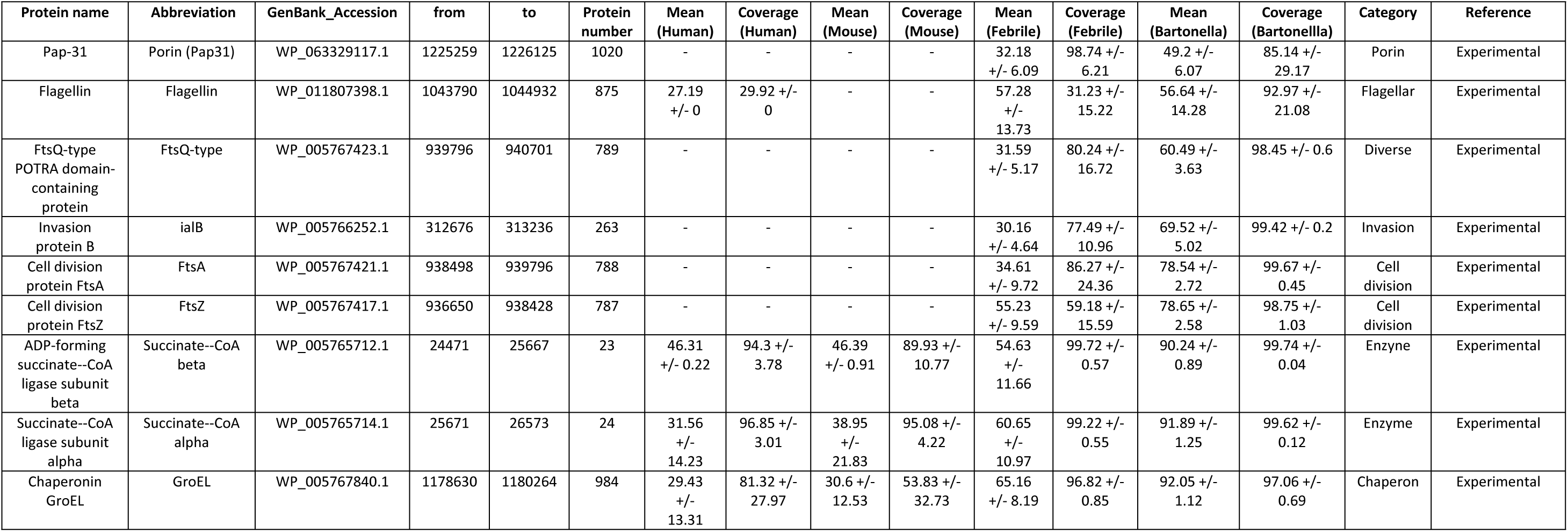
Experimental proteins

### Outer membrane beta-barrel proteins

#### Autotransporter outer membrane

The following 6 autotransporter OMPs (ATOMP), hereafter referred to with capital letters from A to F, were identified: ATOMP-A (WP_005767706.1, 926), ATOMP-B (WP_063329126.1, 949), ATOMP-C (WP_005766622.1, 439), ATOMP-D (WP_005767753.1, 948), ATOMP-E (WP_011807332.1, 438), ATOMP-F (WP_052467648.1, 947), all containing one “autotransporter beta-domain” and one “pertactin domain” with percent identity between 35% and 47% and percent coverage between 50% and 93% (this percentage only in ATOMP-F) for *Bartonella* species other than *B. bacilliformis*. For the beta-barrels, we did not find protein homologs in human, mouse or febrile illness-related species proteomes (Figures 2 and 3). 3D modelling corroborated the beta-barrel structures in these proteins (Figure 4).

**Figure 2.**
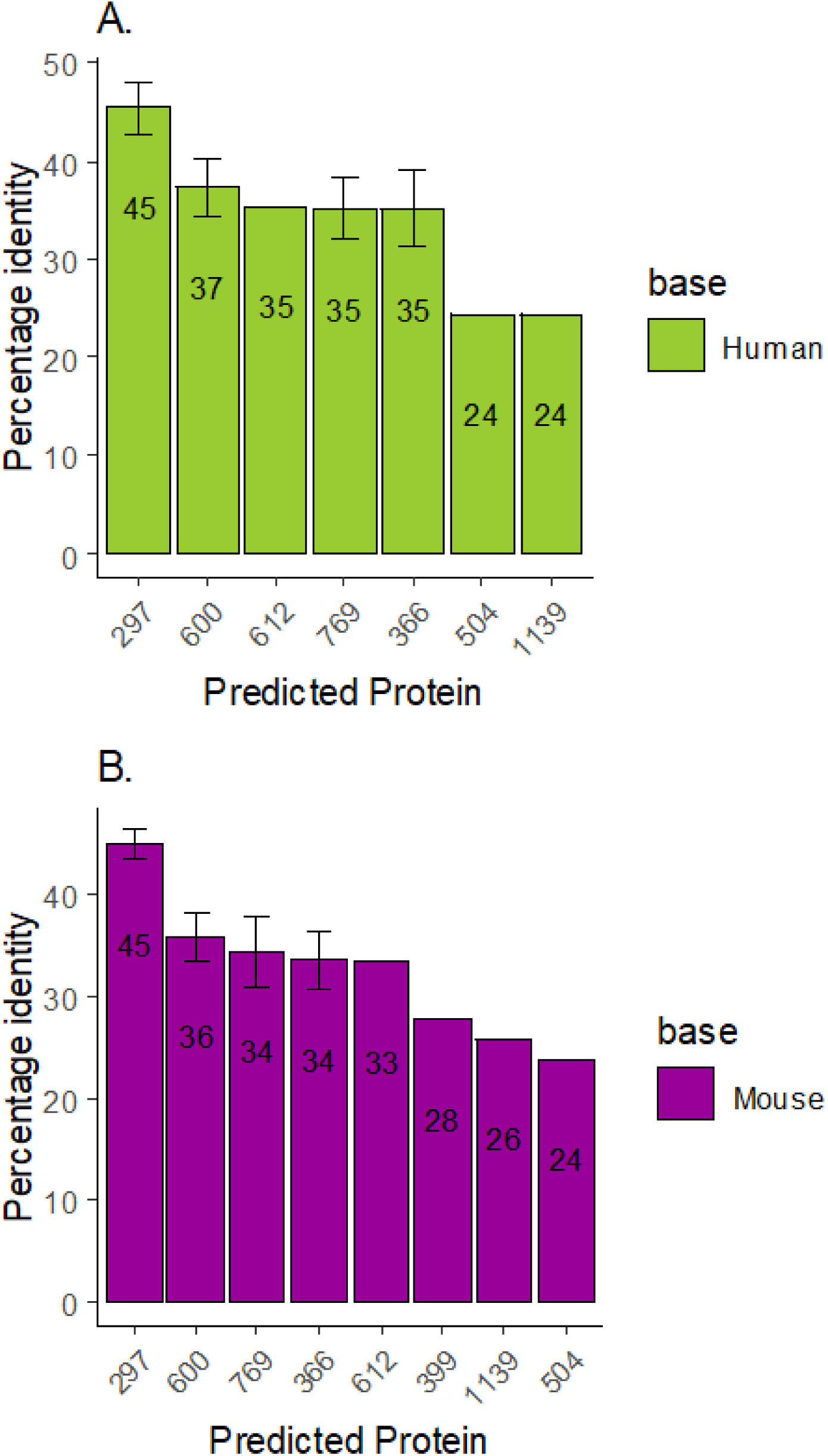
Comparison of outer membrane beta-barrel proteins. Comparison of beta-barrel proteome-proteome in *Bartonella bacilliformis* with *Homo sapiens* (A) and *Mus musculus* (B). X-axis numbers represents the protein number in the annotated reference genome.

**Figure 3.**
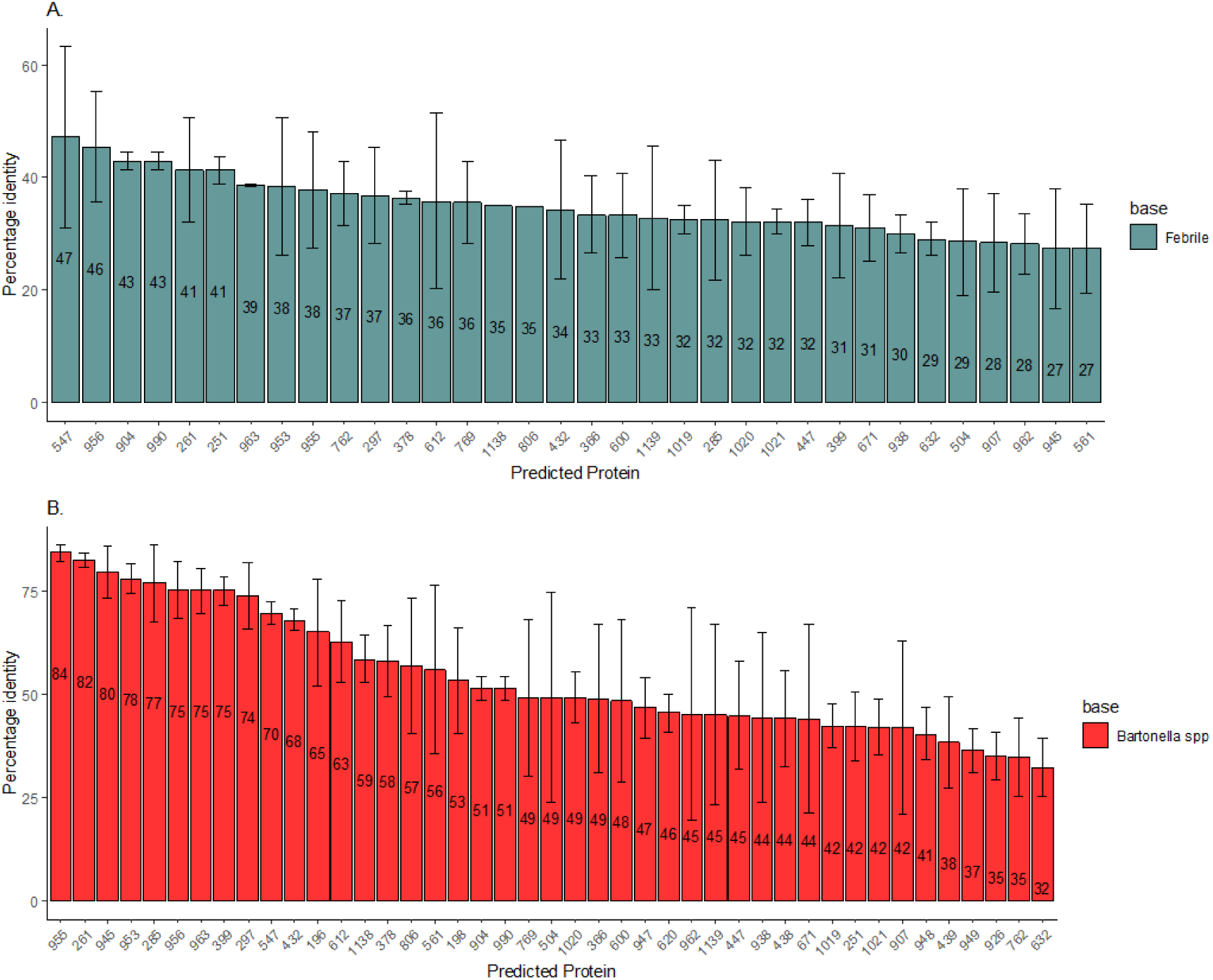
Comparison of outer membrane beta-barrel protein. Comparison of proteome-proteome in *Bartonella bacilliformis* with febrile-associated species (A) and *Bartonella* spp. (B). X-axis numbers represents the protein number in the annotated reference genome.

**Figure 4.**
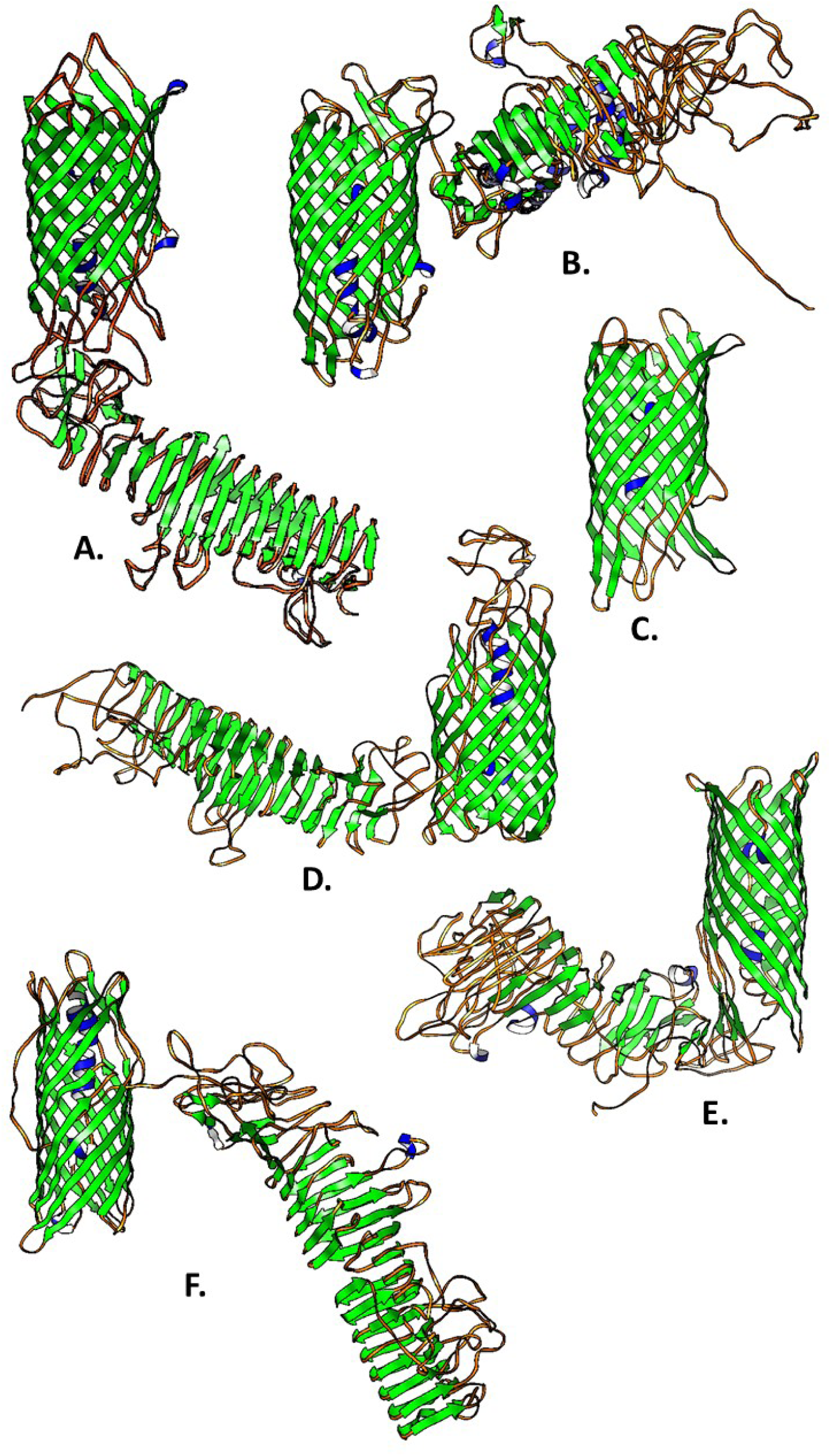
Three dimensional structures of selected outer-membrane beta-barrel: ATOMP-A (A), ATOMP-B (B), Porin-A (C), ATOMP-C (D), ATOMP-D (E) and ATOMP-E (F). secondary structures are colors as beta-strand (green), alpha-helix (blue) and coil (orange). Secondary structures show beta-strand (green), alfa-helix (blue) and coil (orange).

#### Trimeric autotransporter adhesins

Trimeric autotransporter adhesins (TAAS) YadA (WP_005766217.1, 251) were identified, presenting a mean percent similarity of 42.38 +/- 8.33% and a mean percent coverage of 30.12 +/- 21.08% with other *Bartonella* species proteins. When compared with febrile illness-related species proteomes, YadA showed a mean percent similarity of 41.34 +/- 2.46 and mean percent coverage of 30.12 +/- 21.08 (Figure 3).

#### Porins

Four OMBB proteins were identified as porins, three of which are hereafter referred to with capital letters from A to C; these were Porin-A (WP_005767902.1, 1021), Porin-B (WP_005767896.1, 1019), Pap-31 (WP_063329117.1, 1020) and Porin-C (WP_005766494.1, 378) with percent identity between 32% and 36%, percent coverage between 84% and 98% compared with febrile-associated species proteome, and percent identity between 42% and 58% and percent coverage between 83% and 100% compared with *Bartonella* species other than *B. bacilliformis* proteomes (Figure 3). The fourth porin was Pap-31, which is described below in section 3.3.

#### Flagellar

Six beta-barrels were identified as flagellar proteins: FlgE (WP_005767730.1, 938), FlgF (WP_005767784.1, 962), FlgA (WP_005767768.1, 956), FlgH (WP_005767762.1, 953), FliF (WP_005767748.1, 945) and FlgI (WP_005767766.1, 955). However only FlgE and FlgF presented percent similarities below 70% (between 44% and 45% and percent coverage between 48% and 84%) compared to other *Bartonella* proteomes. The remaining proteins, FlgA, FlgH, FliF and FlgI had percent similarities between 75% and 84% and percent coverage between 86% and 97%. All flagellar beta-barrels presented similarities with febrile illness-associated species proteins with percent identities between 30% and 39% and percent coverages between 55% and 91% (Figure 3).

#### Secretory / Efflux proteins

Four proteins involved in secretion and efflux activities were identified: HlyD (WP_005767664.1, 907), TolC (WP_104408431.1,561), TamB (WP_063329116.1, 1138) and signal peptidase I (WP_005766544.1, 399). HlyD, TolC and TamB presented mean percent similarities between 42% and 59%, with percent coverages above 84% compared with other *Bartonella*, whereas the mean percent similarities were less than 35% compared with febrile illness-associated species proteomes. Signal peptidase I showed a mean percent similarity of 75% and a mean percent coverage of 99% (Figure 3).

#### Membrane protein assembly / Translocases

Four proteins involved in membrane proteins assembly or translocation were identified: Omp85 (WP_005768133.1, 1139), BamA (WP_005766757.1, 504), LptD (WP_005766604.1, 432) and BamE (WP_005766845.1, 547). The last two proteins corresponded to lipoproteins and are explained in other sections. Omp85 and BamA presented percent identities between 45% and 50% and percent coverage greater than 82% compared with other *Bartonella*, while percent identities were between 28% and 33% with percent coverage greater than 75% compared with febrile illness-associated species proteomes (Figure 3). They also presented percent similarities between 23% and 26% and percent coverages between 53% and 62% compared with human and mouse proteomes (Figure 2).

### Outer membrane lipoproteins

The present methodology identified 9 outer membrane lipoproteins. Only one lipoprotein “Do family serine endopeptidase” (WP_005767379.1, 769) presented mean percent similarities and mean percent coverages close to 35% with human and mouse proteins (Figure 5). The 3D structures are shown in Figure 6.

**Figure 5.**
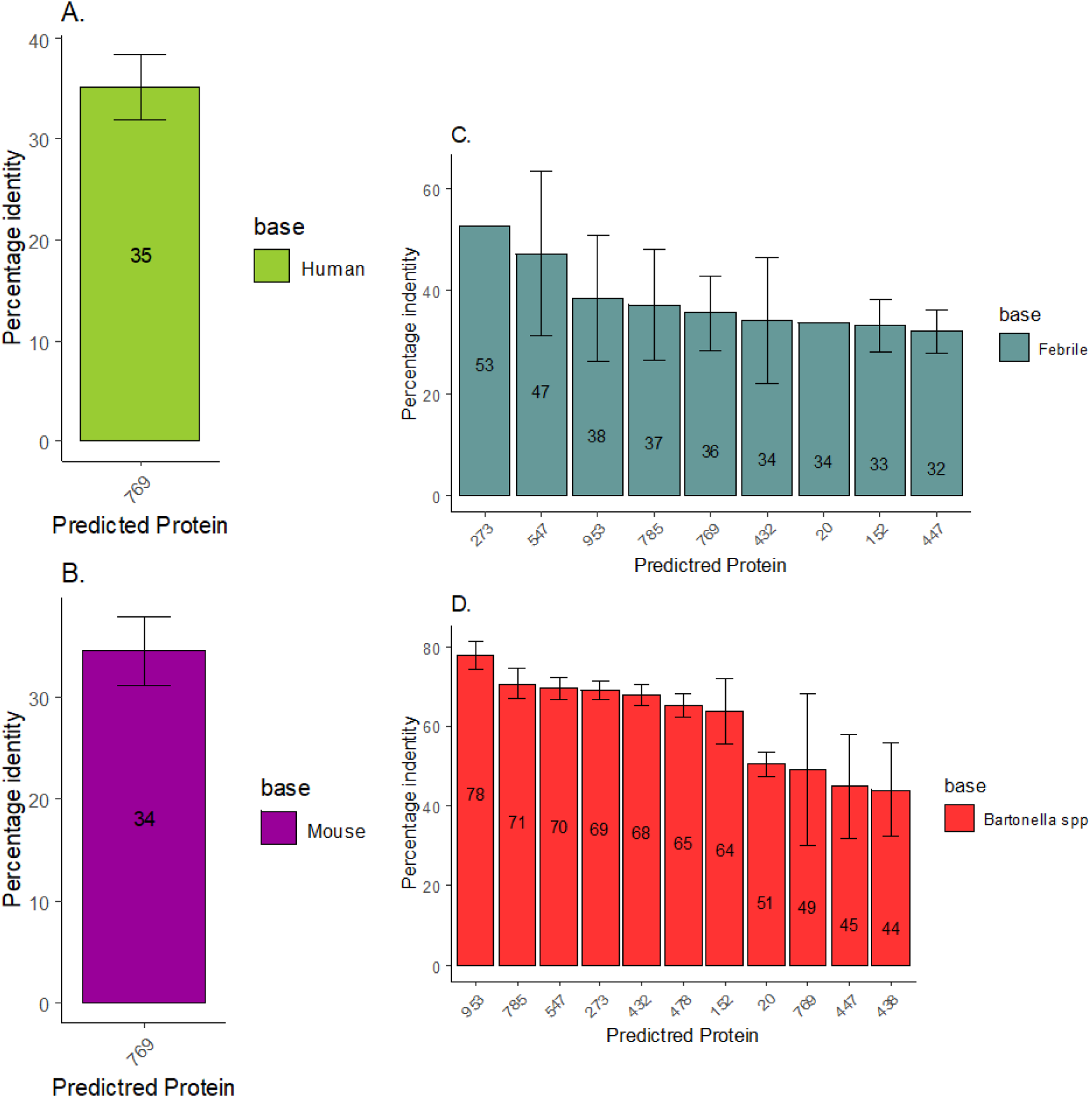
Comparison of outer membrane lipoproteins. Comparison of proteome-proteome in *Bartonella bacilliformis* with *Homo sapiens* (A), *Mus musculus* (B), febrile-associated species (C) and *Bartonella* spp. (D). X-axis numbers represents the protein number in the annotated reference genome.

**Figure 6.**
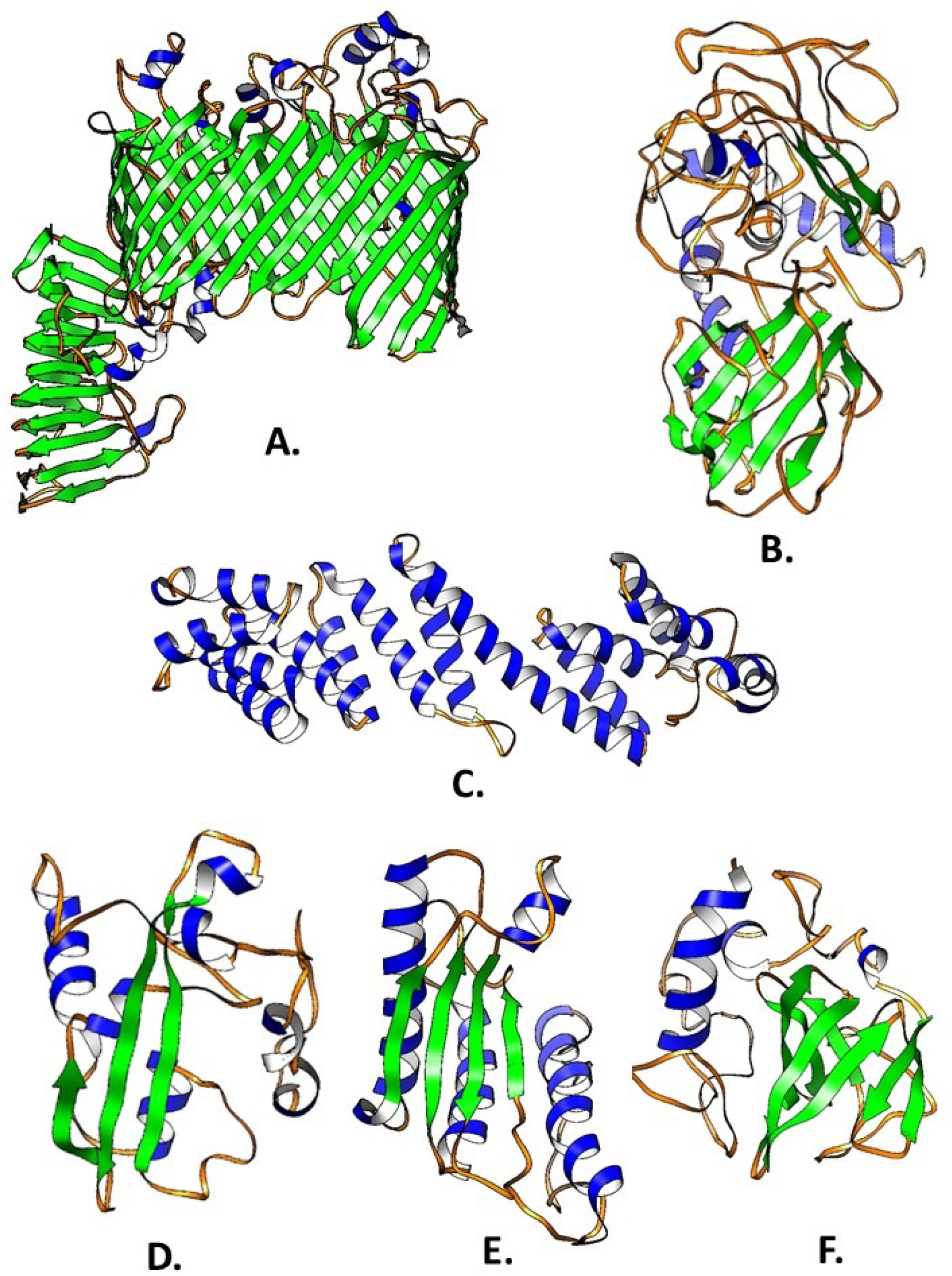
Three dimensional structures of selected outer-membrane lipoproteins: LPS-assembly protein LptD (A), 43kDa - Peptidoglycan DD-metalloendopeptidase (B), BamD (C), BamE (D), Peptidoglycan-associatedlipoprotein-Pal (E) and AprI/Inh family metalloprotease inhibitor (F). Secondary structures are colors as beta-strand (green), alfa-helix (blue) and coil (orange). Secondary structures show beta-strand (green), alfa-helix (blue) and coil (orange).

#### Enzymes/Enzymes regulators

Three lipoproteins involved in enzyme regulations were identified: Peptidoglycan DD-metalloendopeptidase (WP_005766636.1, 447), AprI/Inh family metalloprotease inhibitor (WP_005765706.1, 20) and Do family serine endopeptidase (WP_005767379.1, 769). The first two proteins presented a mean percent identity between 32% and 34%, a mean percent coverage between 36% and 100% in comparison with febrile illness-associated species proteomes, and a mean percent identity between 44% and 51%, and a mean percent coverage between 65% and 100% in comparison with *Bartonella* species proteomes. As described above, only Do family serine endopeptidase presented similarities with human and mouse proteins (Figure 5).

#### Membrane protein assembly / Translocases

We identified 3 lipoproteins involved in protein assembly and translocation activities: lipopolysaccharide (LPS)-assembly protein LptD (WP_005766604.1, 432), OMP assembly factor BamE (WP_005766845.1, 547) and OMP assembly factor BamD (WP_005767413.1, 785). These proteins presented a mean percent identity of 68% to 71%, a mean percent coverage close to 100% in comparison with *Bartonella* species proteomes, and a mean percent identity between 34% and 48% and a mean percent coverage between 75% and 93% compared with febrile illness-associated species proteomes. LptD contains a beta-barrel domain (Figure 6).

#### Outer membrane integrity

The peptidoglycan-associated lipoprotein Pal (WP_005765972.1, 152) was identified as part of membrane integrity, presenting a mean percent identity of 64%, a mean percent coverage of 93% in comparison with *Bartonella* species proteomes, and a mean percent identity of 33% and mean percent coverage of 64% compared with febrile illness-associated species proteomes (Figure 5).

#### Autotransporter outer membrane

One autotransporter outer membrane lipoprotein, hereafter called ATOMP-G (WP_011807332.1, 438) containing also a pertactin domain was identified. This protein presented a mean percent similarity of 44% and percent coverage of 53% only with other *Bartonella* species (Figure 5).

#### Flagellar

One flagellar lipoprotein was identified as the flagellar basal body L-ring protein FlgH (WP_005767762.1, 953). This protein presented a mean percent similarity of 78% and coverage of 98% only with other *Bartonella* species, and a percent identity of 33% and coverage of 65% with febrile illness-associated species proteomes (Figure 5).

### Known antigenic Bartonella bacilliformis proteins

Nine *B. bacilliformis* proteins with known immunological reactivity were also considered. Flagellin (WP_011807398.1, 875), Succinyl-CoA synthetase-β (SCS-β, WP_005765712.1,23), Succinyl-CoA synthetase -α (SCS-α, WP_005765714.1, 24) and GroEL (WP_005767840.1, 984) presented a mean percent identity between 27% and 47% and percent coverages between 29% and 100% compared with human proteins. No similar proteins for flagellin were found compared with those of mouse, but for the three-rest proteins mentioned above, they presented mean percent similarities between 30% and 47% and a mean percent coverage between 53% and 90%. These proteins also presented the highest percent similarities with other *Bartonella* species proteins ranging from 90% to 92% and with a percent coverage ranging from 97% to 100%. Cell division proteins FtsA (WP_005767421.1, 788) and FtsZ (WP_005767417.1, 787) presented mean percent similarities around 78% and percent coverages around 98% with other *Bartonella* species proteins. Three proteins, Porin Pap-31 (WP_063329117.1, 1020), FtsQ-type (WP_005767423.1, 789) and Invasion protein IalB (WP_005766252.1, 263) presented mean percent similarities between 49% and 70% and mean percent coverages between 85% and 99% compared with *Bartonella* species proteins. Porin Pap-31 showed the lowest percent similarity (49.2 +/- 6.07) and percent coverage (85.14 +/- 29.17). Overall, these 9 *B. bacilliformis* proteins presented mean percent similarities between 30% and 65% and a mean percent coverage between 77% and 99%, compared with febrile illness-associated species proteomes (Figure 7).

**Figure 7.**
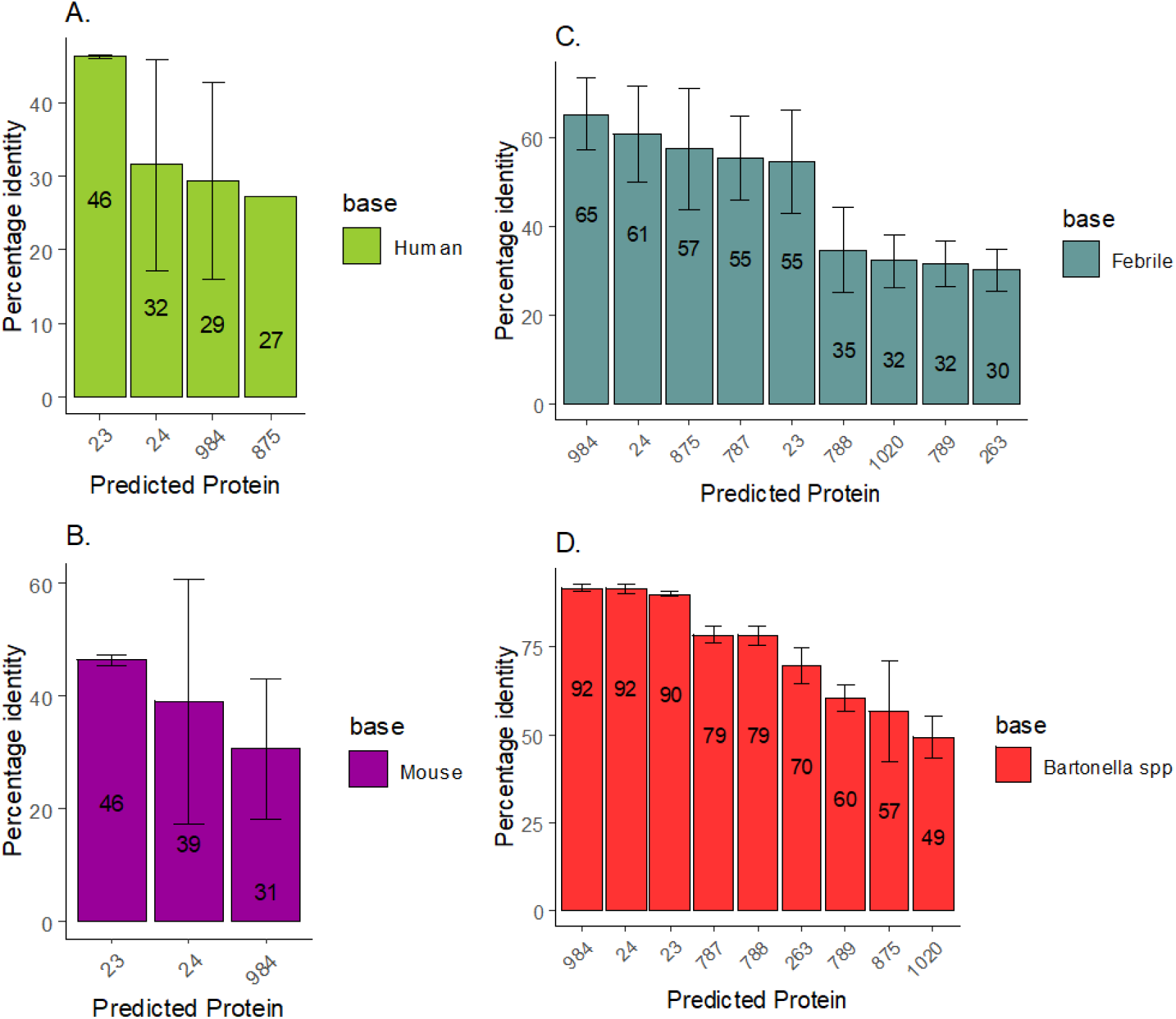
Comparison of known antigenic *Bartonella bacilliformis* proteins. Comparison of proteome-proteome in *Bartonella bacilliformis* with *Homo sapiens* (A), *Mus musculus* (B), febrile-associated species (C) and *Bartonella* spp. (D). X-axis numbers represents the protein number in the annotated reference genome.

## Discussion

In this study we identified OMBB and OMLP of *B. bacilliformis*. Some of these proteins have been identified and or described by other researchers while others have not previously been reported. To achieve the objectives of the study, different OMBB predictors, lipoprotein predictors, subcellular localization predictors and signal-peptides predictors were used. *In-silico* identification of candidate antigenic proteins has been used to propose vaccine-candidates and in the search for antibody targets in the development of serological tests. *B. bacilliformis* outer-membrane and extracellular proteins have been analyzed by the identification of epitopes in the design of an artificial multi-epitope protein with positive results in murine model [37]. In a recently published article, Dichter *et al.* [38] tested the immunogenicity of 22 candidate *B. bacilliformis* proteins identified by a reverse vaccinology approach using Vaxign-1 software. The present study identified 19 exact or homologs of these proteins.

### Outer membrane beta-barrel proteins

#### Autotransporter outer-membrane beta-barrel proteins (ATOMP)

Consist of beta-barrel and pertactin passenger domains, both with immunogenic activity in *B. henselae* [39]. Adhesin and cytotoxin properties have largely been recognized in ATOMP in other microorganisms, such as *Bordetella pertussis* [40], *Shigella flexneri* [41], *Escherichia coli* [42], *Salmonella enterica* [43], *Orientia tsutsugamushi* [44], and *Rickettsia conorii* [45]. The 6 ATOMPs identified in the present study are consistent with the results obtained by Dichter *et al.* (2021) [38] using IgG-ELISA tests. These authors reported one ATOMP (WP_011807413.1) as the most immunodominant antigen and we identified a homolog (WP_063329126.1) of this protein with 43.63% similarity.

#### Trimeric Autotransporter Adhesins

TAAs have been considered as potential candidates for the development of vaccines in several microorganisms, including members of the *Bartonella* genus [46]. The present analysis showed two members of this family: Vomp and YadA-like. Nevertheless, Vomp could not be modeled by Phyre2. These proteins contain 7 and 2 “coiled stalk” domains, respectively, with low percent identities and coverages compared with febrile and other *Bartonella* species.

#### Porins

*B. bacilliformis* porins have been analyzed in different studies as potential antigenic candidates, with Pap31 being the most largely described. Taye *et al.* (2015) described the immunogenicity of Pap-31 and suggested its potential usefulness for the development of rapid diagnostic tolos [47]. The antigenicity of Pap-31 was confirmed by Angkasekwinai *et al.* (2014) [48] and Gomes *et al.* (2016) [10]. Similarly, Pap-31 was detected in the study by Dichter *et al.* (2021) [38](see section 4.3 on Known Proteins). Furthermore, Gomes *et al.* (2021) [11] identified immunogenic peptides within Pap-31 and SCS-α, establishing their inter- and intrageneric discrimination capacity. In agreement with these data, Pap-31 was identified in the present *in-silico* analysis. Furthermore, coinciding with Dichter *et al.* (2021) [38], 3 other porins (A, B and C) were identified. Interestingly, these porins present one of the lowest percent similarities with other *Bartonella* species (48%-58%). In this sense, *B. bacilliformis* and B. ancashensis Pap-31 contains amino acid tandem repeat motifs possibly implicated in immune evasion because of its high antigenicity and disorder structure, which is absent in the remaining genomes of established *Bartonella* spp. [49].

#### Flagellar proteins

It has been suggested that the release of LPSs through outer membrane vesicles containing virulence factors is mediated by flagellar activity [50]. Immunogenic, adhesive and invasive properties have been identified in flagellar proteins of other species, such as *Brucella abortus* FlgJ [51], *Borrelia burgdorferi* FlgE [52], *Pseudomonas aeruginosa* FlgE [53] and *Clostridium difficile* FliC and FliD [54]. Of the 6 flagellar proteins identified in the present study, flagellar hook FlgE and flagellar basal body L-ring FlgH were also reported by Dichter *et al.* (2021) [38], whereas we could not find references for flagellar basal-body rod FlgF, flagellar basal body P-ring formation FlgA, flagellar M-ring FliF and flagellar basal body P-ring protein FlgI. Nevertheless, percent similarities close to (or greater than) 78% compared with other *Bartonella* species limit the use of these proteins as discriminant candidates but not as vaccine or generic targets.

#### Secretory-Efflux

As in other cases, data about *Bartonella* secretory proteins are scarce or absent. In other species, such as *B. pertussis*, *Listeria monocytogenes* or *Streptococcus sanguinis*, among others, the involvement of members of this protein group, such as SipX, SipZ or CyaD (homologous to HlyD), has been observed in virulence processes, including intracellular multiplication [55–57]. While no data about HlyD, TolC, signal peptidase-I and the DUF490 domain of *B. bacilliformis* have been found, these proteins are probably involved in the pathogenic process of *B. bacilliformis*, and might thereby qualify as antigen candidates despite a percent similarity greater than 70%.

#### Membrane protein assembly / Translocases

In Gram-negative bacteria the Bam complex is responsible for OMP assembly and maintenance [58]. The Bam complex consists of BamA to BamE proteins, and BamD and BamE are lipoproteins. In other species these proteins have been considered as promising immunogenic proteins [59–63]. While, in the present case, these proteins might be considered as antigenic candidates of *B. bacilliformis*, the similarity of close to 70% with other *Bartonella* of the 6 detected proteins belonging to this category suggests they may play a better role as genus markers. However, caution should be taken considering that Omp85-TamA and BamA share 25% similarity with human and mouse proteins and thereby being at risk of presenting false positive results.

### Outer Membrane lipoproteins

#### Enzymes/Enzymes regulators

Of the 5 members of this group detected, the Peptidoglycan DD-metallo-endopeptidase was tested in western blot assays [64]. In agreement with our results, further studies considered this protein as a good antigenic candidate in the serological diagnosis of CD [65]. While there are no data on the remaining proteins detected as belonging to this group, by inference with what has been described in other microorganisms, including members of the genus *Bartonella*, they might be involved in virulence processes [66–68].

#### Membrane protein assembly/Translocases

The LptD protein is involved in the assembly of polysaccharides in the outer membrane [69]. Mass spectrometry analysis conducted by Li *et al.* (2014) [70] in immunized mice-derived-sera detected LptD as a promising immunogenic protein against *Vibrio parahaemolyticus*, and in a vaccine assay performed in 2016, Zha *et al.* [71] distinguished 100% immune protection suggesting its use as a vaccine and for antibody-based therapy. In the present case, the subcellular localization of LptD could favor its use as an antigenic protein in *B. bacilliformis*. However, previous studies have described a low antigenicity [38].

#### Outer membrane integrity

Peptidoglycan-associated lipoprotein (Pal) is involved in bacterial pathogenicity, being present in more than 100 species [72]. For instance, studies conducted in *Haemophilus ducreyi* related the pathogenesis of chancroid to a mutant version of the encoding gene Pal, highlighting the importance of Pal as a virulence factor [73]. Meanwhile, peptidyl-prolyl isomerase, which has been involved in correct protein folding and pathogenicity processes [74, has been identified as highly antigenic in *Acinetobacter baumannii* [75]. Both proteins represent or interact with virulence factors in different species. Despite the lack of data for *B. bacilliformis*, a similar role for these proteins cannot be ruled out. The similarities between 68% and 78% with other *Bartonella* species and the non-negligible similarities of peptidyl-prolyl isomerase with human and mouse proteins alerts about their use as therapeutic targets.

#### Domains with unknown function (DUF)

The DUF490 domain is present in the translocation and assembly module protein (TamB) involved in the secretion of virulence-associated OMPs in intracellular bacterial species such as *B. burgdorferi* in conjunction with BamA [76]. It is, thus, possible that the unique DUF490 domain-containing protein identified in *B. bacilliformis* could play a similar role. The two different DUF1561 domain-containing proteins of unknown function identified in the present study have both similar and low identities (58%) with those of other *Bartonella* species. Previous studies on *B. bacilliformis* also identified homologs of these proteins as immunogenic [38].

#### Tail fiber domain-containing protein

One tail fiber protein, that is present in many, if not all, *Bartonella* species was detected in the present study. Tail fiber proteins are present in bacteriophages and mediate viral-host invasion [77, 78].

### Known proteins

The immunogenicity of *B. bacilliformis* flagellin, Brps (TAAs), IalB, FtsZ, 43kDa-lipoprotein (LppB) and Pap31has been reviewed suggesting the inclusion of these antigen candidates in vaccine strategies [79]. The high levels of identities for SCS-β, SCS-α and GroEL with human, mouse and febrile-associated species proteins makes them inadequate for the serological diagnosis of *B. bacilliformis*, but they might be useful for generic detection of *Bartonella* spp. On the other hand, according to the present search, Pap-31 represents the best candidate. presenting a percent similarity close to 50% with other *Bartonella* species. Although it has been detected in previous studies focused on the search for antigenic candidates [10,47,48], Dichter *et al.* [38] observed a low reactivity, with only two of 26 patient samples showing IgG positive results, which were attributed to conformational differences. While these results disagree with the results of Taye *et al.* (2005) [47] and Angkasekwinai *et al.* (2014) [48], they agree with those obtained by [10] Gomes *et al.* (2016) who did not observe IgG reactivity of Pap31 and with the reported presence of a series of number variable amino acid tandem repeats [49], resulting in both sequence differences between *B. bacilliformis* isolates, as well as the presence of a disordered region, usually leading to a non-stable conformation. These findings alerting about the use of Pap31 for the detection of IgG. The amino acid tandem repeat region is placed just behind the 16 amino acid peptide 187-QAIGSAILKGTKDTGT-202 showing high IgM immunogenic levels [11, 49]. While this peptide is conserved in all genomes (with a single amino acid difference in the case of the strain Ver097 (RefSeq: WP_041848867), the above mentioned Pap31 data highlight the need to establish its usefulness as a marker of acute infections in serum of patients with infections related to *B. bacilliformis* isolates presenting differences in the amino acid tandem repeat region.

#### Perspectives and implications of this study

Adequate medical interventions rely on disease diagnosis. CD includes an acute phase, characterized by anemia and febrile illness and a chronic phase distinguished by skin eruptions [2]. Furthermore, the presence of asymptomatic carriers has been reported, being considered as reservoirs and perpetuators of the disease [2, 10]. It is therefore important for a serological test for CD to not only be sensitive and specific, but it should also be able to detect early (acute) and late (chronical) infections, as well as asymptomatic carriers. Detection of the latter is of special interest to advance towards the eradication of CD [2]. The development and implementation of IgM (acute phase) and IgG (chronic phase and previously exposed population) antibody tests, respectively is important, with ELISA, Immunoblot and immunochromatographic lateral flow being good alternatives. The validation of these methods should include the geographic representation of CD from different endemic areas of Ecuador and Peru.

In addition to their use for diagnostic purposes, many of the *B. bacilliformis* proteins identified could act as therapeutic targets (inhibiting the secretion of virulence proteins, inhibiting the maintenance of cell wall) or vaccines (by designing a mix of complete antigens, multiepitope proteins, or outer-membrane vesicles containing virulence factors). Future studies should consider both 1) the genetic variability of *B. bacilliformis*, and 2) the genetic variability of the human immune components (HLA-I, HLA-II) and should be carried out using molecular dynamics approaches.

## Conclusion

We report the *in-silico* identification of OMBB and OMLP in *Bartonella bacilliformis*, exploring a combination of predictors at the genome level. The list of 32 beta-barrel proteins and 9 lipoproteins identified in this study could be used for the development of target therapies, serological diagnostic tests and vaccine candidates for the control of CD.

## Acknowledgments

This work was supported by Fondo Nacional de Desarrollo Científico, Tecnológico y de Innovación Tecnológica (FONDECYT) presented to Giovanna Mendoza-Mujica as part of the project “Evaluación de candidatos antigénicos recombinantes para la innovación del diagnóstico confirmatorio de la Enfermedad de Carrión” (E041-2019-01, register number: 63531).

We thank Donna Pringle for language correction.

